# Development of a genetically encoded and potent PDE6D inhibitor

**DOI:** 10.1101/2025.09.29.679187

**Authors:** Atanasio Gómez-Mulas, Elisabeth Schaffner-Reckinger, Hanne Peeters, Rohan Chippalkatti, Arnela Dautbasic, Matthew James Smith, Shehab Ismail, Daniel Kwaku Abankwa

## Abstract

PDE6D is a trafficking chaperone of prenylated proteins, such as small GTPases. Several small molecule inhibitors have been developed against it, given that the oncogene K-Ras is one of the cargo proteins. Inhibitor development suffered from the fact that inhibitors against the hydrophobic pocket of PDE6D were typically poorly water-soluble. Here we describe the development of genetically encoded inhibitors that are inspired by high-affinity natural cargo of PDE6D. Our most potent inhibitor, SNAP-STI, encodes merely a farnesylated tetra-peptide, which efficiently blocks PDE6D binding of farnesylated cargo. Direct comparison with small molecule PDE6D inhibitors suggests its higher potency. We show that inhibition of K-Ras membrane anchorage and K-RasG12C-dependent MAPK-signaling by SNAP-STI is weak, consistent with what is observed after PDE6D knockdown. Our data therefore further support that PDE6D is not a suitable surrogate target for efficient inhibition of K-Ras membrane anchorage and MAPK-activity. Nonetheless, by exploiting contacts at the pocket entry, we established a generalizable strategy to design high-affinity PDE6D inhibitors, providing powerful tools for PDE6D biology and target validation.

## Introduction

The Ras-MAPK-pathway regulates vital cellular processes, including proliferation and differentiation. Its overactivation is associated with cancer and developmental diseases collectively called RASopathies, which affect multiple organs ^[1]^. GTP-loading switches the conformation of Ras, which initiates downstream signaling by recruiting effector proteins to Ras at the plasma membrane. For example, recruitment of the effector Raf to membrane-associated Ras initiates the MAPK-pathway ^[2]^. Farnesylation of the CAAX-box of the hypervariable region provides the affinity of Ras proteins towards membranes. In this motif, the cysteine (C) is prenylated, followed by proteolysis of the AAX-tripeptide (aliphatic residue A and any residue X) and carboxymethylation. Long-range diffusion of Ras through the cytosol requires trafficking chaperones, which shield the prenyl moiety. This is necessary to feed the active vesicular transport of Ras to the plasma membrane ^[3]^.

The trafficking chaperone PDE6D has been proposed as a surrogate target for K-Ras4B (hereafter K-Ras) ^[4]^. It binds the farnesyl-membrane anchor at the C-terminus of K-Ras, but also other prenylated proteins, including dually prenylated small GTPases, including Rab1B, Rab4A and Rab7A ^[5]^. PDE6D is also a major trafficking chaperone of prenylated proteins operating in the primary cilium, such as the lipid phosphatase INPP5E ^[6]^. The primary cilium is an antenna-like membrane protrusion on most stem-/progenitor-cells where it serves as a hub for several developmental-signaling pathways, such as Wnt and Hedgehog ^[7]^. The distinct localization of PDE6D cargo to the bulk plasma membrane and primary cilium is mediated by the two small GTPases Arl2 and Arl3, respectively. When GTP-bound, they bind to PDE6D to release low-affinity cargo, such as K-Ras (GTP-Arl2) or high-affinity cargo, such as INPP5E, in addition to low-affinity cargo (GTP-Arl3) ^[6, 8]^.

The fact that Ras plasma membrane localization is necessary for its activity led to the development of farnesyltransferase inhibitors (FTI) as the first Ras drug targeting approach. These inhibitors failed in the clinic, due to alternative prenylation of K-Ras and N-Ras by geranyl-geranyltransferase I, the two Ras isoforms most frequently mutated in cancer ^[9]^. Given that PDE6D facilitates plasma membrane localization of K-Ras ^[10]^, a number of small molecule inhibitors were developed against PDE6D ^[9a]^, including our own Deltaflexin1, -2 and -3 ^[11]^. Deltaflexin3 is a highly water-soluble, low nanomolar inhibitor of the prenyl-binding pocket of PDE6D. However, during its development, we also noticed that compounds with a higher affinity become less soluble and acquire more off-target activities ^[11b]^. Like with other PDE6D inhibitors, its ability to shut down MAPK-signaling was surprisingly low considering it indirectly blocks K-Ras signaling. However, this low activity was consistent with the fact that PDE6D directs only about 25-50 % of the plasma membrane trafficking of K-Ras ^[11b]^. Moreover, we noticed discrepancies with the reported target affinity values. It is common to establish the in vitro affinity of inhibitors to PDE6D in a competitive fluorescence polarization assay using fluorescently labelled atorvastatin as a PDE6D-binding probe ^[4]^. However, when measuring the affinity with a Rheb-derived farnesylated peptide or in an SPR-assay directly looking at the interaction between K-Ras and PDE6D, only micromolar affinities were recovered ^[11]^.

A way forward in PDE6D inhibitor development could be inspired by its natural cargo, where affinity is modulated not only by contacts within the hydrophobic prenyl-binding pocket of PDE6D but also by interactions at the pocket entrance. For example, the protein INPP5E binds with K_D_ = 4 nM to PDE6D as compared to K_D_ = 2.5 μM for binding of K-Ras ^[6, 12]^. This almost 1000-fold increase is mediated by only two residues at position -1 and -3 upstream of the prenylated cysteine ^[5-6]^.

## Results and Discussion

We aimed at generating genetically encoded, high-affinity binders to PDE6D by exploiting contacts at the entrance of the prenyl-binding pocket of PDE6D. (**Figure 1A**). This approach was inspired by the nanomolar binding affinity of INPP5E that is mediated by two residues at the -3 and -1 position relative to the prenylated cysteine (**Figure 1A**). Mutating these two lysine residues of K-Ras to the serine and isoleucine (SI) found at the corresponding positions of INPP5E, resulted in the K-Ras-SI mutant ^[5]^.

**Figure 1.**
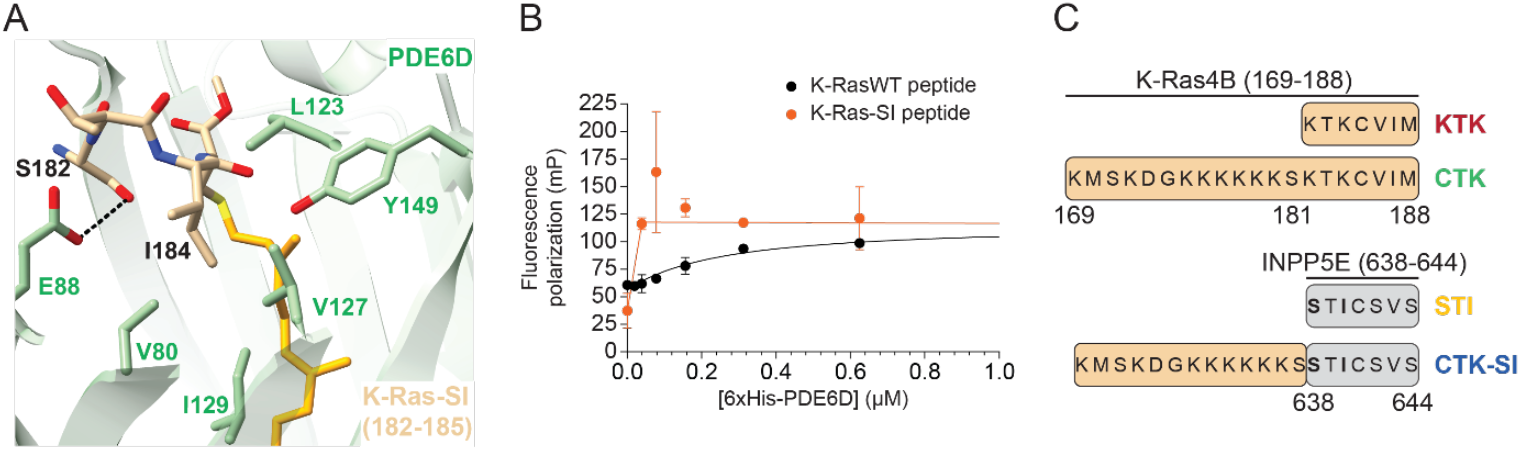
Genetically encoded peptides derived from INPP5E are high-affinity PDE6D binders. **A**) Crystal structure of K-Ras-SI (only residues 182-185 are shown in tan, farnesyl moiety in orange) and PDE6D (green) (PDB ID: 7Q9U). H-bond between the side chain of Ser182 and Glu88 of PDE6D is indicated by dashed lines. **B**) Fluorescence polarization binding data of 50 nM wild type K-Ras-derived peptide (FAM-KKKKKKSKTKC-Far-OMe) and K-Ras-SI-derived peptide (fluorescein-DGKKKKKKSSTIC-Far-OMe) and increasing concentrations of 6xHis-PDE6D. Means ± SD are plotted of n = 3 or n = 2, respectively, independent biological repeats. Solid lines represent the quadratic binding equation fit (K-RasWT peptide) and the two linear equation fits consistent with an active site saturation titration regime (K-Ras-SI peptide). **C**) Names of the genetically encoded PDE6D binders and their sequence composition relative to K-Ras (tan) and INPP5E (grey).

Fluorescence polarization assays confirmed binding of a farnesylated and carboxymethylated peptide derived from the C-terminus of wild type (WT) K-Ras to PDE6D with a K_D_ = 0.20 ± μM (**Figure 1B**), consistent with prior reports ^[13]^. By contrast, a K-Ras-SI-derived peptide reached saturation rapidly when 39 nM of PDE6D were titrated against 50 nM of the peptide, indicating an affinity far exceeding that of the wild type (**Figure 1B**).

BRET-biosensors can assay the effective binding or competitive activity of genetically encoded inhibitors in HEK 293 (hereafter HEK) cells ^[14]^. Plate reader-based BRET experiments can detect binding events inside intact cells following a distance-dependent (< 12 nm) energy transfer between tagged proteins ^[15]^. One protein of interest is tagged with a Renilla Luciferase variant (e.g., RLuc8 or NanoLuc, nL), which catalyzes the enzymatic conversion of the chemical substrate coelenterazine 400a to provide the donor energy. Energy is transferred to a close acceptor-fluorophore (e.g. GFP2 or mNeonGreen, mNG) that is genetically fused to an interaction partner. Using PDE6D and K-Ras-derived BRET pairs, we confirmed that the high increase in affinity observed in vitro (**Figure 1B**) was maintained in the interaction-dependent BRET of the K-Ras-SI mutant (**Figure S1A**).

We reasoned that the last seven amino acids of INPP5E (STI-CSVS, with C representing the prenylated cysteine), represent a minimal sequence for a genetically encoded high-affinity PDE6D inhibitor. To this end, we genetically fused this sequence via a 15-residue-long GS-linker to the C-terminus of a SNAP-tag. In addition, we generated a longer hybrid version where the seven INPP5E residues were N-terminally extended by residues 169-181 of K-Ras (CTK-SI). The matching controls were the corresponding peptides named KTK and CTK, which were entirely derived from K-Ras (**Figure 1C**). Note that CTK-variants describe the 20-mer polypeptide derived from the K-Ras C-terminus, where the designation ‘CTK’ has been established for many years ^[16]^.

We evaluated these four PDE6D binders in a BRET assay designed to measure the displacement of K-Ras from PDE6D. Unlike with full-length K-Ras-SI (**Figure S1B**), no clear potency differences were apparent between the four SNAP-tagged constructs (**Figure S2A**). Swapping of the donor and acceptor pair did not alter the results with BRET values remaining low, suggesting a low fraction of the proteins was interacting (**Figure S2B**). We reasoned that the low BRET signal and the difficulty resolving potency differences likely reflect the small cytosolic pool of K-Ras available to bind PDE6D, which limits the dynamic range of the biosensor.

Indeed, the subcellular distribution of analogous EGFP-variants of the four unique binders varied considerably in C2C12 myoblasts, with EGFP-CTK showing the maximum plasma membrane localization and EGFP-STI the lowest (**Figure 2A**). In line with the SI-mutation increasing affinity to PDE6D and thus solubilization, co-expression of PDE6D sequestered EGFP-STI and EGFP-CTK-SI to the nucleo-cytoplasm (**Figure 2B**). Nuclear localization is consistent with the small size of the PDE6D/EGFP-binder construct complexes.

**Figure 2.**
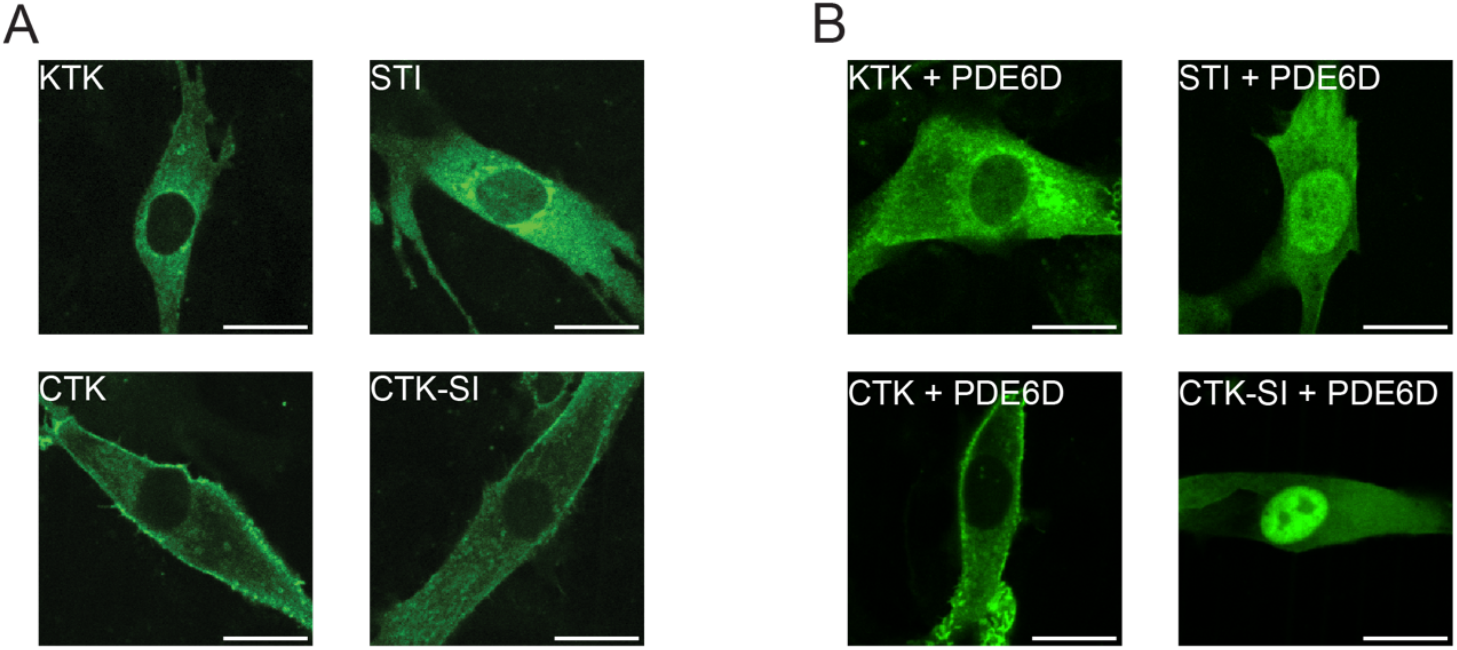
Genetically encoded PDE6D binders show distinct subcellular distributions. **A,B**) Confocal images of C2C12 myoblasts transfected with the EGFP-tagged variants of PDE6D binding sequences (A), or cotransfected with a 1:1 plasmid ratio of EGFP-tagged PDE6D binders and mCherry-PDE6D (B). Scale bar = 20 µm.

To improve the signal and dynamic range of our PDE6D biosensor, we therefore employed a KTK-derivative as BRET-acceptor, which displayed a higher cytosolic fraction able to engage with PDE6D (**Figure 3A**). This modified BRET-biosensor allowed a clear discrimination of PDE6D inhibitory activity, which decreased in the order STI, CTK-SI>>KTK>CTK (**Figure 3A**). In line with this, direct PDE6D binding in BRET-experiments with the acceptor-tagged PDE6D binders followed the same order STI, CTK-SI>>KTK>CTK (**Figure S3A**).

**Figure 3.**
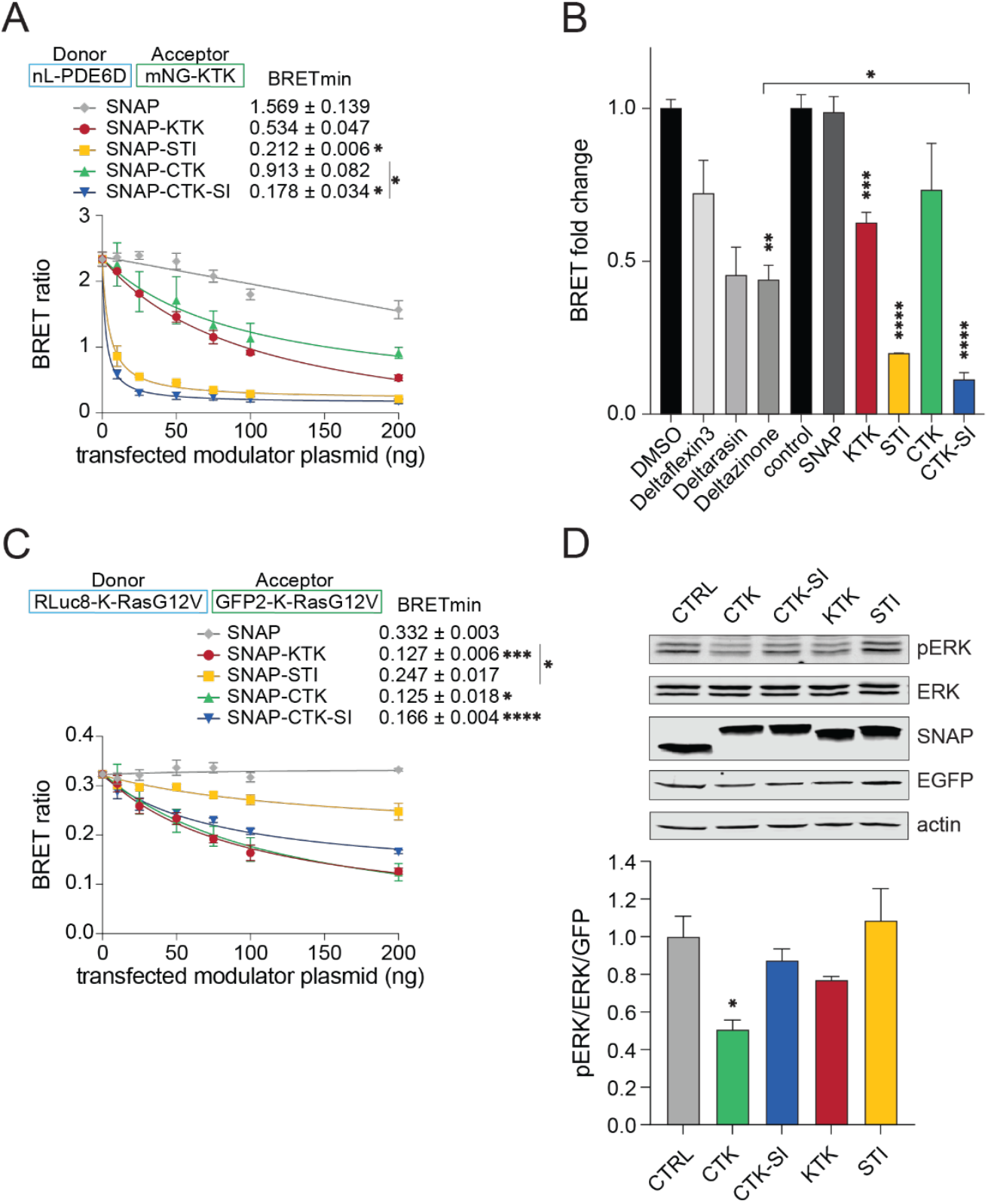
SNAP-STI appears to inhibit PDE6D stronger than small molecule inhibitors. Dose-dependent inhibitory effect of SNAP-tagged PDE6D binders on nL-PDE6D/mNG-KTK BRET (donor-acceptor plasmid ratio was 1:10); n = 3. **B)** Normalized BRET ratio of nL-PDE6D/mNG-KTK BRET experiments testing at 20 µM Deltaflexin3, Deltarasin or Deltazinone and SNAP-tagged PDE6D binders (50 ng of transfected plasmid, normalized to 0 ng of transfected modulator control); n = 3. **C**) Dose-dependent inhibitory effect of SNAP-tagged constructs on RLuc8-K-RasG12V/GFP2-K-RasG12V BRET (donor-acceptor plasmid ratio was 1:10); n ≥ 3. **D**) Representative immunoblots of the phosphorylation of ERK1/2 in HEK cells transfected with mEGFP-K-Ras-G12C and genetically encoded SNAP-tagged PDE6D binders. Antibodies used for labeling are indicated. The plot shows the quantification of relative ERK phosphorylation; n = 4. Statistical analysis as compared to the control condition was performed with the Kruskal-Wallis test.

To compare the inhibitory activity of our genetic constructs with small molecule PDE6D inhibitors, we prepared a calibration curve to estimate the approximate cytosolic concentration of SNAP-STI and SNAP-KTK per HEK cell (**Figure S3B**). We determined that 50 ng of transfected DNA yielded an intracellular concentration of ∼10 μM. Direct comparison with three small molecule PDE6D inhibitors: Deltaflexin3, Deltarasin, and Deltazinone tested at 20 μM, suggested that SNAP-STI and SNAP-CTK-SI are more potent at inhibiting PDE6D ^[4, 11b, 17]^ (**Figure 3B**).

Given that the subcellular distribution of some inhibitors was similar to that of K-Ras with partial localization to the plasma membrane (**Figure 2A**), we assessed disruption of K-Ras nanoclustering. Nanoclusters are di-/oligomeric proteo-lipid assemblies of Ras, which are necessary for its signaling and represent a potential drug targeting opportunity ^[18]^.

Using an established K-Ras-nanoclustering BRET assay in the absence of PDE6D overexpression ^[19]^, we found that inhibitors disrupted K-Ras nanoclustering in line with their plasma membrane abundance in the order CTK, KTK>CTK-SI>STI (**Figure 3C**). A decrease in the K-RasG12V/K-RasG12V nanoclustering-dependent BRET signal may also indicate an overall drop in functional K-Ras membrane anchorage, as we demonstrated previously ^[11b, 20]^. At low acceptor-to-donor plasmid ratios, the same order was found for the proximity BRET of mNG-derivatives with K-Ras (**Figure S3C**). However, at a higher ratio, CTK-SI behaved more similarly to CTK, probably because endogenous PDE6D became saturated with CTK-SI and could no longer sequester it, leaving it to distribute similarly to parental CTK.

Finally, inhibition of MAPK-signaling in HEK cells (CTK>KTK, CTK-SI>STI) (**Figure 3D**) followed the order of how inhibitors effectively impacted K-Ras membrane anchorage (**Figure 3C**), rather than PDE6D inhibition (**Figure 3A**).

These data therefore support that disruption of K-Ras nanoclustering is more potent than inhibition of PDE6D for MAPK-activity suppression. Similarly, when comparing the impact of the inhibition of K-Ras prenylation and thus plasma membrane anchorage using the mevalonate pathway inhibitor mevastatin, or a combination of farnesyl transferase and geranylgeranyl transferase inhibitors, with that of PDE6D-ablation we observed a significant >60 % drop in pERK-levels with the former treatment (**Figure S3D**), but only a ∼20 % drop with nearly complete PDE6D knock-down (**Figure S3E**). Altogether, these data underscore that PDE6D is not a suitable surrogate target of K-Ras-MAPK-signaling.

Our results suggest that it is possible to raise high-affinity PDE6D inhibitors by exploiting contacts at the entrance of the prenyl-binding pocket.

We developed SNAP-STI, a bio-relevant inhibitor that can serve as a benchmark for the cellular and in vivo assessment of small molecule PDE6D inhibitors. This will be important to resolve, for instance, if broad inhibition of PDE6D is toxic in the adult, as loss of PDE6D during development leads to the severe developmental disease Joubert-Syndrome ^[21]^. Given that the final inhibitor is essentially a farnesylated tetrapeptide, it is plausible that a peptidomimetic with a similar potency can be developed. Indeed, others have synthesized a laurylated 11-mer peptide as an inhibitor of UNC119, which is highly related to PDE6D ^[22]^. Novel applications of PDE6D inhibition could thus be explored, which may include modulation of ciliary cargo proteins and the activity of the cilium to regulate stem cell function ^[23]^.

Our results suggest that a natural cargo-inspired affinity increase of inhibitors of PDE6D or its related trafficking chaperones has untapped potential.

## Acknowledgements

This work was supported by the Luxembourg National Research Fund (FNR) grants AFR/23/17112420/Bil_ABANKWA_SPREDCanUL2 to DKA and MJS, INTER/FWO/23/18086068 molGluRAS2 to DKA, and FWO grant G042824N to SI. MJS holds a Canada Research Chair (CRC) in Cancer Signalling and Structural Biology. We thank Zoe Geimer for technical support.

## Experimental Methods

### Peptides

Wild-type K-Ras peptide (FAM-KKKKKKSKTKC-OMe-Far) was synthesized by Biosynth and K-Ras-SI derived peptide (fluorescein-DGKKKKKKSSTIC-OMe-Far) was synthesized by JPT Innovative Peptide Solutions.

### Plasmid cloning

Expression constructs were prepared using gateway recombination cloning technology (Thermo Fisher Scientific). To this end, we used either gBlock gene fragments with the cDNA of the genetically encoded inhibitors (Integrated DNA Technologies), or plasmids with cDNA constructs from the Ras-Initiative (K-Ras4BG12V, K-Ras4BG12C from kit#1000000089, and PDE6D from #R702-E30), all inserts flanked by BP cloning sites. Using BP Clonase II enzyme mix, the fragments were inserted into pDONR221 plasmids to obtain entry clones flanked by LR cloning sites. Then, using the LR Clonase II enzyme mix, we combined entry clones containing the promoter, N-terminal tag, and cDNA of interest. The resulting pDest-305 plasmids were amplified in the ccdB-sensitive *E*.*coli* strain DH10B and validated by Sanger sequencing.

### Cell lines

C2C12, HEK293 c18 (both from ATCC), and HEK293-EBNA cells (gift of Florian M. Wurm, EPFL, Lausanne, Switzerland) were cultured in Dulbecco’s modified Eagle’s medium (DMEM) containing 2 mM L-glutamine, 1 % penicillin/streptomycin, and ∼9 % (v/v) fetal bovine serum (high serum). Cells were incubated in a humidified atmosphere with 5 % CO_2_ at 37 °C.

### Protein purification

6xHis-PDE6D was purified using a protocol adapted from a previously published protocol ^[5]^. A His6-tagged PDE6D construct in a pRSF-Duet vector was transformed into BL21 DE3 *E. coli* cells. Following the transformation, precultures were grown at 37°C overnight and transferred to large cultures at 37°C until OD600 nm reached 0.5. The bacteria were then induced with 0.2 mM isopropyl β-D-1-thiogalactopyranoside (IPTG) and protein was expressed for 16 h. The bacteria were harvested by centrifugation and stored at -80°C until use. Bacterial pellets were thawed and resuspended in lysis buffer containing 50 mM Tris pH7.5, 300 mM NaCl and 2 mM Dithiotreitol (DTT) with chicken hen lysozyme before lysis using a microfluidizer at 20,000 psi. The bacterial lysate was then centrifuged at 18,600 rpm and supernatants was loaded onto a 5 mL HisTrap column (Cytiva) at 5 mL/min. The column was then washed with 24 mM imidazole and protein was eluted using a gradient of 24-300 mM imidazole in a 50 mM Tris pH7.5, 150 mM NaCl and 0.5 mM TCEP buffer. The eluate was then passed through a Superdex HiLoad 16/600 75 pg column equilibrated with 50 mM Tris pH7.5, 150 mM NaCl and 0.5 mM TCEP buffer at 0.7 mL/min. 6xHis-PDE6D was then concentrated and snap-frozen in liquid nitrogen before storage at -80°C.

### Fluorescence polarization peptide binding assay

Fluorescence polarization measurements were recorded on a Tecan Spark plate-reader using 96-well Corning flat black half-area non-binding plates at 22°C and using an excitation wavelength of 496 nm and emission wavelength of 524 nm for the K-Ras-derived peptide and a 498 nm excitation wavelength and 527 nm emission wavelength for the K-Ras-SI-derived peptide. All measurements were recorded in a buffer containing 50 mM Tris pH7.5, 150 mM NaCl and 0.5 mM TCEP. Fluorescence polarization was measured following a 30 min incubation of peptides at 50 nM with increasing concentrations of 6xHis-PDE6D. For the binding curve of the K-Ras-derived peptide to PDE6D, the dissociation constant was obtained by fitting the data to a quadratic equation using GraphPad Prism: FP = Fmin − (Fmin − Fmax) * (E + L + Kd-sqrt[(E + L + Kd))^2 − 4*E*L)]/(2*E). FP is the fluorescence polarization signal, Fmin and Fmax is the minimum and maximum polarization signal, E is the K-Ras peptide concentration, L is the 6xHis-PDE6D concentration and K_d_ is the dissociation constant as a measure for affinity. For the active site titration of K-Ras-SI-derived peptide to PDE6D, the linear regimes were fitted with a linear equation Y=slope*L+intercept.

### Bioluminescence Resonance Energy Transfer (BRET)

BRET experiments were conducted as previously described ^[14-15]^. Briefly, 220,000 HEK 293-EBNA cells were seeded per well in 12-well plates, transfected after 24 hours with the plasmid constructs using 2 µL of JetPrime, and measured 48 h later. Donor saturation titration BRET was performed as previously described by us. From these data we eventually determined the pseudolinear regime for the optimal acceptor-to-donor ratio in dose-response experiments. These latter experiements were conducted by adding increasing amounts of a modulator (plasmid or compound) 24 h after transfection, measuring 16 h later. BRET was measured using a CLARIOstar Microplate Reader (BMG Labtech) using white flat-bottom 96-well plates as described ^[14-15]^.

### Confocal microscopy

C2C12 cells were seeded in 6-well plates with 0.17 mm coverslips at 200,000 cells per well. After 24 h, the cells were transfected with JetPrime, and 4 hours later, fresh high serum medium was added to the cells. 48 h after transfection, the cells were fixed with 4 % (w/v) paraformaldehyde in PBS and permeabilized with 0.5 % Triton X-100. The nuclei were stained with 0.2 μg/mL Hoechst 33342 in PBS containing 0.05 % v/v Tween 20 and then washed. Vectashield mounting medium was added to glass slides and the coverslips were mounted on them. The fixed cells on coverslips were imaged using a 60 × NA 1.3 oil immersion objective on a Nikon Ti-E microscope equipped with a Yokogawa CSU-W1 spinning disk confocal unit and an Andor iXon Ultra EMCCD camera. The EGFP-fluorophore was detected by 488 nm laser excitation and 535/20 nm band pass filter detection. Z-stacks were acquired with 0.6 μm spacing. Images were acquired with Nikon NIS-elements software and analyzed in Fiji/ImageJ.

### Immunoblotting

Prior to transient transfection, 100,000 HEK293 c18 cells were seeded in 2 mL DMEM/well in 6-well plates. After 24 h, cells were transfected with 100 nM siRNA as reported before ^[11b]^, and 3.75 μL Lipofectamine RNAiMAX and after 48 h, transfection was performed with 0.5 μg of each plasmid DNA and 3 μL Lipofectamine 2000. In situ cell lysis was performed 24 h after plasmid transfection as described before in ice-cold lysis buffer (50 mM Tris-HCl pH 7.5, 150 mM NaCl, 0.1 % w/v SDS, 5 mM EDTA, 1 % v/v Nonidet P-40, 1 % v/v Triton X-100, 1 % w/v sodium-deoxycholate, 1 mM Na_3_VO_4_, 10 mM NaF, 100 μM leupeptin) containing cocktails of protease inhibitors and phosphatase inhibitors ^[11b]^. After clarification of the lysates by centrifugation, the total protein concentration was determined by performing a Bradford assay. A standard curve with bovine serum albumin was established. SDS-PAGE was performed using 10 % w/v homemade polyacrylamide gels to resolve proteins (40 μg per lane) under reducing conditions. Proteins were then transferred by semi-dry transfer onto nitrocellulose membranes. Saturation was performed for 1 h at 22 °C in PBS containing 2 % w/v BSA and 0.2 v/v % TWEEN 20, and membranes were incubated overnight at 4 °C with primary antibodies diluted in saturation buffer. Incubation with corresponding secondary antibodies diluted in saturation buffer was carried out for 1 h at 22 °C. At least three wash steps in PBS 0.2 % TWEEN 20 were performed after each antibody incubation. For each blot, *β*-actin levels were determined as a loading control. An Odyssey Infrared Image System (LI-COR Biosciences) was used to quantify signal intensities. ERK phosphorylation was calculated as described before ^[11b]^, and data were scaled to the corresponding control present on each blot.

### Estimation of the expression of SNAP-tagged peptides in cells

On immunoblots, band intensities corresponding to SNAP-tagged peptides encoded by the PDE6D binder constructs were determined using an anti-SNAP antibody and were expressed as ratios vs. a known amount of purified SNAP protein (New England Biolabs) loaded on the same gel. Considering the approximate number of cells loaded per well and using an approximate HEK293-EBNA cell volume of 2.5 pL ^[24]^, a dose response curve of the concentration of SNAP-tagged peptide per cell vs. the amount of transfected DNA was established. Using Graph Pad Prism software, simple linear regression was performed to obtain the linear fit of the dose-response curve, and the R^2^ value was determined.

### Statistical analysis

Graph Pad Prism software was used to analyze data. Unless otherwise indicated, data plots show mean values ± SEM. The number of independent biological repeats is indicated by n. A Brown-Forsythe and Welch ANOVA analysis was performed comparing the BRET ratio value of the highest modulator amount (BRETmin) between all samples, or the highest A/D plasmid ratio values (BRETtop) between all samples, unless stated otherwise in the legends. A p-value < 0.05 was considered statistically significant, with significance levels annotated as: \**p* ≤ 0.05; \*\**p* ≤ 0.01; \*\*\**p* ≤ 0.001; \*\*\*\**p* ≤ 0.0001.

