## Supplementary material for "Development of a genetically encoded and potent PDE6D inhibitor": Supplmental Information - Figures

### Supporting Information Gómez et al.

#### Supplementary Figures

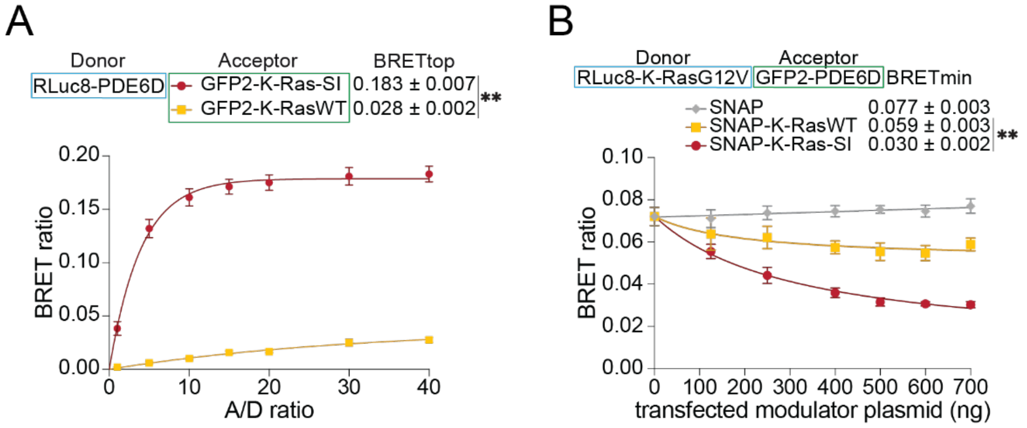


**Figure S1. Data supplementing main Figure 1.**

**A**) BRET donor saturation titration curves of 25 ng RLuc8-PDE6D donor plasmid titrated with increasing amounts of acceptor plasmid GFP2-K-RasWT, or GFP2-K-Ras-SI. A/D ratio gives the acceptor-to-donor plasmid ratio; n ≥ 3. BRETtop values were analyzed with a two-tailed Welch's t-test. **B**) Dose-dependent effect of the SNAP-tagged PDE6D binders on RLuc8-K-RasG12V/ GFP2-PDE6D BRET (donor-acceptor plasmid ratio = 1:10); n ≥ 3.

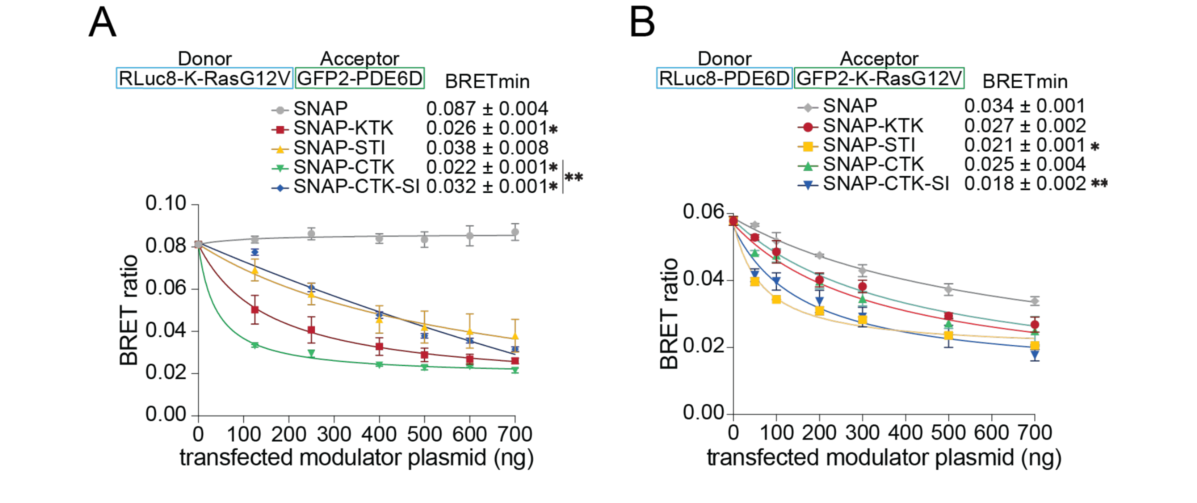


**Figure S2. Data supplementing main Figure 2.**

**A, B**) Dose-dependent effect of the SNAP-tagged PDE6D binders on RLuc8-K-RasG12V/ GFP2-PDE6D BRET (donor-acceptor plasmid ratio = 1:10); n ≥ 3 (A), or on RLuc8-PDE6D/ GFP2-K-RasG12V BRET (donor-acceptor plasmid ratio = 1:20); n ≥ 3 (B).

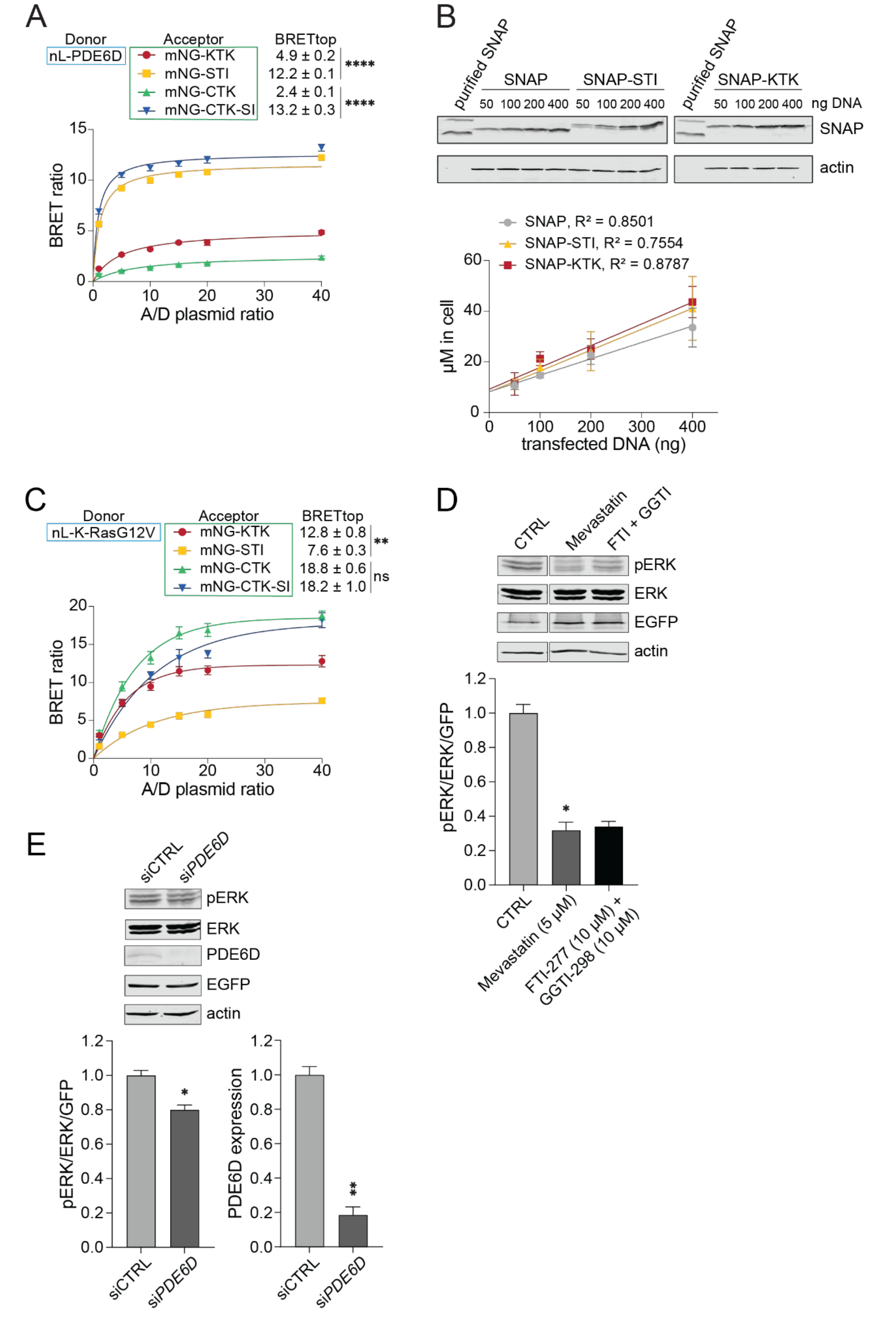


**Figure S3. Data supplementing main Figure 3.**

**A**) BRET donor saturation titration curves of 25 ng nL-PDE6D with increasing amounts of the mNG-tagged PDE6D binder constructs; n = 5. **B**) Representative immunoblots showing the expression of SNAP-tagged PDE6D binders and 2.5 μg purified SNAP protein loaded on the same gel. SNAP corresponds to the lower band in the lane of purified SNAP. Antibodies used for labeling are indicated. The calibration curves show the estimated concentration of SNAP-tagged constructs per cell vs. the amount of transfected DNA. Mean ± SD are shown. A simple linear regression was fitted to the data, and the $R^2$ value was determined. **C**) BRET donor saturation titration curves of 25 ng nL-K-RasG12V donor plasmid with increasing amounts of the mNG-tagged PDE6D binder constructs; n = 5. **D, E**) Immunoblot analysis of the phosphorylation of ERK1/2 (pERK) in HEK cells transfected with EGFP-K-Ras-G12C and treated overnight with Mevastatin or a combination of farnesyltransferase inhibitor FTI-277 and geranyl-geranyltransferase inhibitor GGTI-298 (D) or following siRNA-mediated PDE6D downregulation (E). Antibodies used for labeling are indicated in representative immunoblots. The plots show the quantification of relative ERK phosphorylation or PDE6D expression; n = 4. Statistical analysis as compared to the control condition was performed with the Kruskal-Wallis test (D) or the Mann-Whitney test (E).
